# Cortical circuit principles predict patterns of trauma induced tauopathy in humans

**DOI:** 10.1101/2024.05.02.592271

**Authors:** Helen Barbas, Miguel Angel Garcia-Cabezas, Yohan John, Julied Bautista, Ann McKee, Basilis Zikopoulos

## Abstract

Connections in the cortex of diverse mammalian species are predicted reliably by the Structural Model for direction of pathways and signal processing (reviewed in ^1,2^). The model is rooted in the universal principle of cortical systematic variation in laminar structure and has been supported widely for connection patterns in animals but has not yet been tested for humans. Here, in *postmortem* brains of individuals neuropathologically diagnosed with chronic traumatic encephalopathy (CTE) we studied whether the hyperphosphorylated tau (p-tau) pathology parallels connection sequence in time by circuit mechanisms. CTE is a progressive p-tau pathology that begins focally in perivascular sites in sulcal depths of the neocortex (stages I-II) and later involves the medial temporal lobe (MTL) in stages III-IV. We provide novel quantitative evidence that the p-tau pathology in MTL A28 and nearby sites in CTE stage III closely follows the graded laminar patterns seen in homologous cortico-cortical connections in non-human primates. The Structural Model successfully predicted the laminar distribution of the p-tau neurofibrillary tangles and neurites and their density, based on the relative laminar (dis)similarity between the cortical origin (seed) and each connection site. The findings were validated for generalizability by a computational progression model. By contrast, the early focal perivascular pathology in the sulcal depths followed local columnar connectivity rules. These findings support the general applicability of a theoretical model to unravel the direction and progression of p-tau pathology in human neurodegeneration via a cortico-cortical mechanism. Cortical pathways converging on medial MTL help explain the progressive spread of p-tau pathology from focal cortical sites in early CTE to widespread lateral MTL areas and beyond in later disease stages.

## Introduction

Cortico-cortical connections are summarized succinctly by the rules of the Structural Model (reviewed in ^1,3^), which uniquely predicts the laminar distribution and, by extension, the probable processing direction in the cortex ^2^. The model is rooted in the classical theory of cortical systematic variation, traced from the phylogenetically ancient limbic areas characterized by the simplest lamination, to areas with progressive elaboration of laminar structure seen through six-layer eulaminate cortices. The classical principle of cortical systematic variation was discovered independently by several investigators working on different continents and with different species (reviewed in ^4^). Recent studies using detailed transcriptomic analyses have described patterns that enrich the set of markers that depict systematic changes in cortical architecture ^5,6^, suggesting a common evolutionary origin of the cerebral cortex (reviewed in ^7^).

The Structural Model relates the laminar pattern of connections to the laminar structural relationship of each pair of linked cortices. Connections that originate in a cortex with higher laminar structure originate in the upper layers and their axons terminate in the middle layers (deep 3-5a) of the target area, in a pattern commonly called feedforward. In the reverse direction, pathways originate in the deep layers 5 and 6 and innervate layer 1, in a pattern called feedback. The model is thus relational so that connections between areas that differ markedly in laminar structure involve fewer layers and are sparse ^8^. In contrast, connections between areas that have comparable laminar structure are balanced within layers and are dense. The key predictions of the relational model uniquely reveal the direction of processing, while density of connections alone is uninformative for direction (^8^; reviewed in ^1^).

Unlike animal studies, demonstration of the patterns of cortical connections in humans with *postmortem* use of tracers has encountered methodological limitations so that labeling mostly involved short-range pathways (e.g., ^9,10^). Here, we used a different approach by study of the laminar pattern of tau pathology in a human neurodegenerative disease, CTE, in which the tau pathology is thought to spread along connected networks via trans-synaptic transport between affected neurons. However, the circuit mechanism of spread in p-tau pathologies has remained elusive ^11–15^.

We used brains from individuals with neuropathologically verified CTE, a neurodegenerative p-tau pathology associated with repetitive mild head trauma, including concussive and nonconcussive head impacts. CTE is defined pathologically by the perivascular accumulation of p-tau as neurofibrillary tangles and neurites preferentially at the sulcal depths of the cortex ^16,17^. The severity of CTE pathology can be divided into a 4-tiered staging scheme, based on the density and regional distribution of p-tau pathology ^16,18^.

The goal of this study was to determine whether the laminar pattern of p-tau pathology in CTE, and thus direction of spread, is consistent with cortico-cortical connection rules, and to simultaneously address the broad question of whether laminar connections in humans can be predicted by a theoretical model. We found that the laminar patterns and density of p-tau pathology in MTL in stage III CTE bear a striking resemblance to patterns of cortico-cortical connections of non-human primates. The graded laminar pattern of p-tau pathology in MTL in CTE follows the rules of the theoretical model and is supported by a computational progression model. Further, we provide evidence that the initial p-tau pathology in the depths of sulci of CTE stages I-II follows a different pattern, based on connectivity within narrow columns. Collectively, these findings provide support for the use of a theoretical model to predict the direction of spread and progression of p-tau pathology over time, as well as patterns of connection in the human cortex.

## Results

### The laminar structure of MTL areas varies systematically from a medial to lateral direction

As in other cortical regions, the MTL shows graded changes in lamination ^19–21^. From a medial to lateral direction, A28 is agranular, and is abutted laterally by dysgranular cortex and then by six-layer eulaminate areas that show laminar elaboration in successive lateral sites of MTL. The architecture of MTL allowed straightforward analysis of p-tau pathology in neurons in single sections without the need to group areas into types ^8^, in a region where areas are interconnected in macaque monkeys ^22,23^.

### The laminar distribution of neurons with p-tau pathology in MTL

Systematic p-tau pathology in the cortex emerges first in entorhinal A28 of MTL in CTE stage III ^17,24^ (Fig. 1a). A28 also has the densest p-tau pathology within MTL (Fig. 1b-k), and thus is akin to the origin or seed site (Fig. 1a, box b). As in cortico-cortical connections, pathways of p-tau transmission appear bidirectional. Accordingly, just as all areas that receive connections from A28 (Fig. 1b) reciprocate, we find that as p-tau from A28 is transmitted to other MTL areas, then these MTL areas reciprocate. The labeled neurons in Fig. 1 (magnified panels below the cross section in a) show the origin of the reciprocal pathways from different MTL sites that project to A28. According to model circuit rules, the relative laminar distribution of neurons that reciprocate projections varies by the difference/similarity in laminar structure between the linked areas (areas with label and the seed site). The smaller the difference in structure, the more evenly neurons are distributed in the upper layers (2-3) and in the deep layers (5-6), as seen in Fig. 1, panels c and d. In more lateral sites, in panels e-k, the laminar distribution of neurons with p-tau pathology shifts gradually to a preponderance in the upper layers. The graded laminar distribution of neurons with p-tau pathology reflects the increasing difference in laminar structure between A28 and sites from e to k, where the few labeled neurons are found increasingly in the upper layers.

**Fig. 1:**
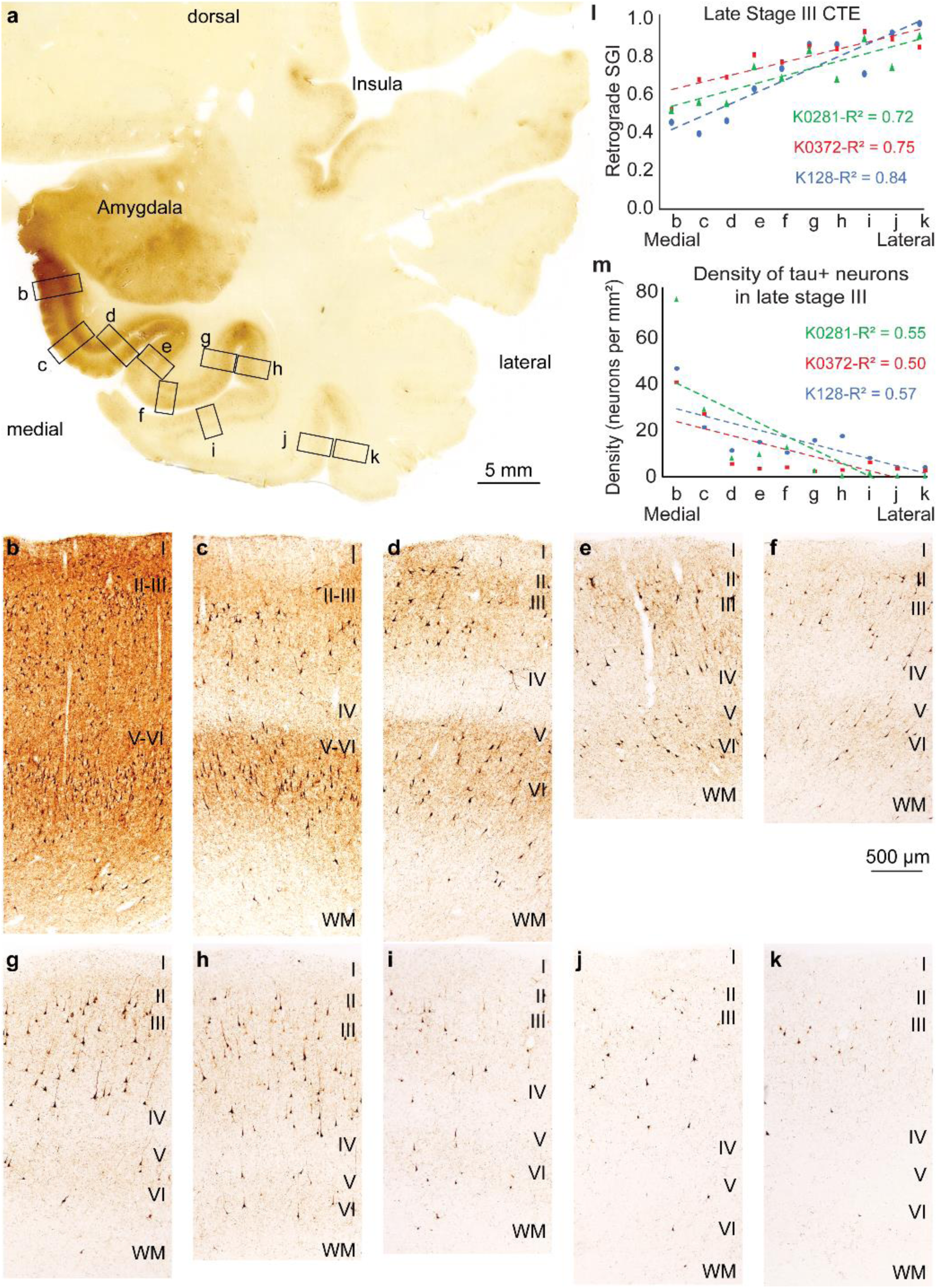
P-tau pathology in temporal cortices in late stage III CTE. **a**, Cross section through the medial temporal lobe (case K128) at the level of the amygdala immunostained against hyperphosphorylated tau (box b shows the densest p-tau pathology in A28). **b-k**, At high magnification, regions (outlined in a) show a graded pattern in the laminar distribution of neurons with p-tau pathology that parallels the laminar connectivity between the agranular A28 (b) through adjacent dysgranular and progressively with eulaminate (six-layer) lateral temporal areas. **l**, Quantitative analysis shows a monotonic increase of neurons with p-tau pathology in the upper layers (SGI) from medial to lateral parts of MTL for three cases with late stage III CTE. **m**, Quantitative analysis of the density of neurons with p-tau pathology in the supragranular layers of three cases with late stage III CTE, gradually decreasing from A28 (b) to progressively adjacent lateral sites within MTL.

The pattern of p-tau pathology was corroborated in two additional cases of stage III CTE. Quantitative analysis showed monotonic increase of neurons with p-tau pathology in the upper layers from medial to lateral parts of MTL (*p = 0.0002, F = 41.07, df = 9.* Fig. 1l (for individual maps and combined analyses in the additional cases see Supplementary Figs. 1-3). The number of neurons was normalized for each of the ten sites and expressed as the supragranular index (SGI), which is the ratio of neurons in the upper layers over all neurons with p-tau pathology at each site. There is about an equal number of neurons with p-tau pathology in the upper and deep layers in medial sites (c-d), with an increasing ratio of neurons in the upper layers in progressively adjacent lateral sites within MTL. The trend in the density of neurons with p-tau pathology is seen with quantitative analysis by using the SGI in Figure 1m for the three late-stage III cases (and Supplementary Fig 3b for combined density: (p = 0.011684, F = 10.57, df = 9).

In another case, we saw the same trend of label with p-tau. Remarkably, this case, which was classified by refined staging as early stage III, revealed a crucial laminar pattern: p-tau pathology in A28 (the seed site) was found predominantly in the deep layers (Fig. 2a, b), unlike advanced stage III, where all layers are affected (Fig. 1b). This finding reveals that p-tau pathology spreads from A28 to lateral MTL sites via feedback pathways. As in the cases above, the laminar distribution pattern and density of label were graded from medial to lateral MTL sites (Fig. 2b-k), and by quantitative analyses (Fig. 2l, m: (*p = 0.000029, F = 71.39, df = 9*).

**Fig. 2:**
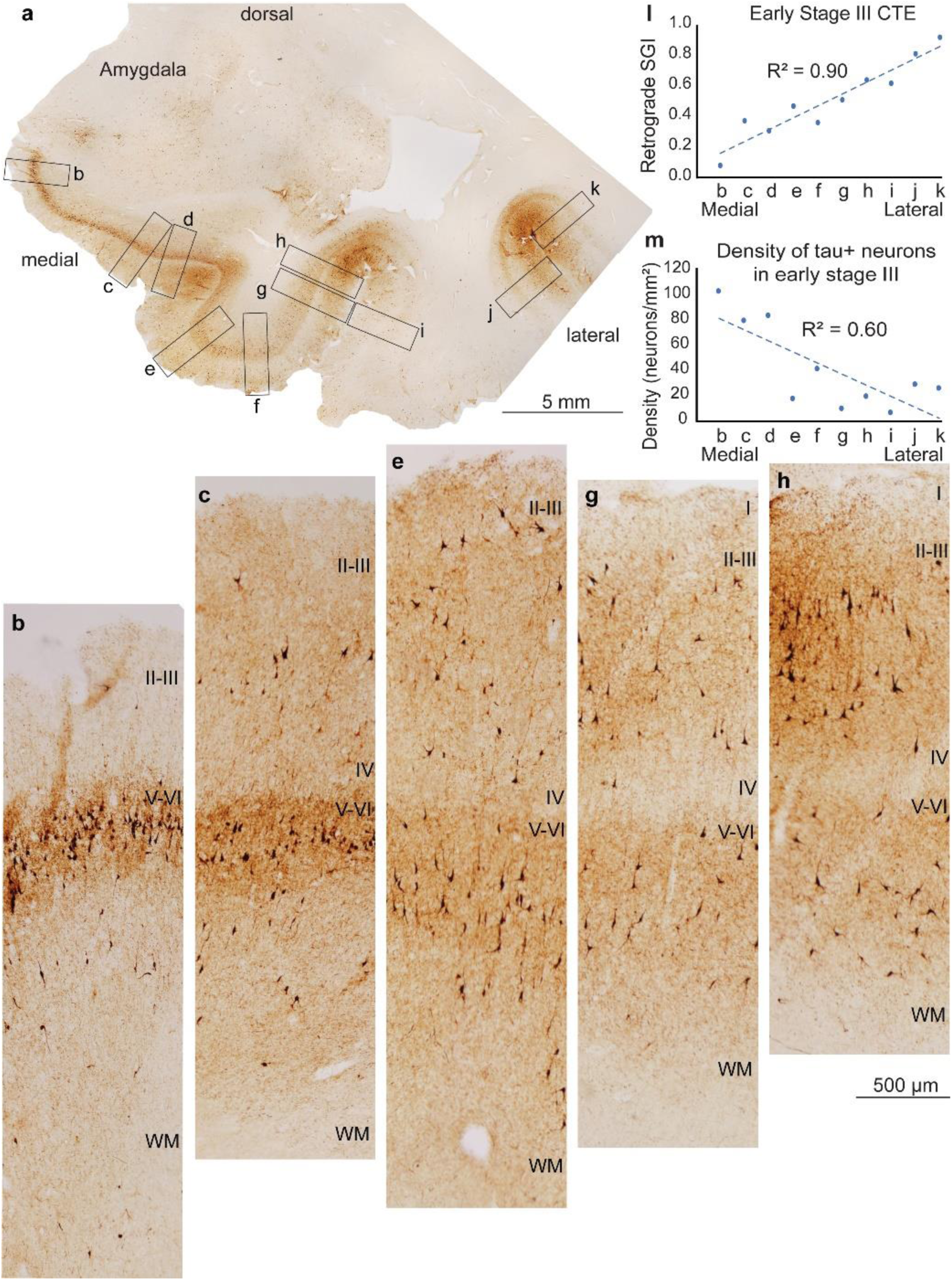
P-tau pathology in MTL cortices in early stage III CTE. **a**, Overview of cross section through the medial temporal lobe at the level of the amygdala (case AT8-CV-SLI-67) immunostained against hyperphosphorylated tau. **b-h**, Sites shown at high magnification (boxes outlined in a) show a graded pattern in the laminar distribution of neurons with p-tau pathology that parallels patterns of laminar connectivity between A28 and areas with increasing laminar differentiation in the MTL. **l**, Quantitative analysis shows monotonic increase of neurons with p-tau pathology in the upper layers (SGI) from medial to lateral parts of MTL for a case with early stage III CTE. **m**, Quantitative analysis of the density of neurons with p-tau pathology in the supragranular layers of a case with early stage III CTE, gradually decreasing from A28 to progressively adjacent lateral sites within MTL.

### The termination of pathways from seed A28 to MTL sites

From the seed site in A28, feedback-like pathways innervate other MTL areas. Antibodies for p-tau pathology label both neurons and axons and their terminals, which allowed us to address whether p-tau pathology in A28 is reflected in the pattern of termination of pathways within MTL areas. This analysis included the density of the fine ‘background’ label in each panel, made up of axon fibers and terminations but excluded labeled neurons (see Methods). We normalized the findings from each site as the SGI, as above. As seen in Figure 3, there is a significant correlation in the SGI within the MTL sites, which is low at sites closest to the seed A28 and increasingly higher in lateral panels (Fig. 3a: *p = 0.0028, F = 18.04, df = 9*; Fig. 3b: *p = 0.012, F = 10.64, df = 9*). This relationship is based on the predominant ‘feedback’ pathways that A28 issues, which terminate increasingly in the upper layers in areas that differ more and more in laminar structure from the seed site in A28 through sequential lateral MTL sites.

**Fig. 3:**
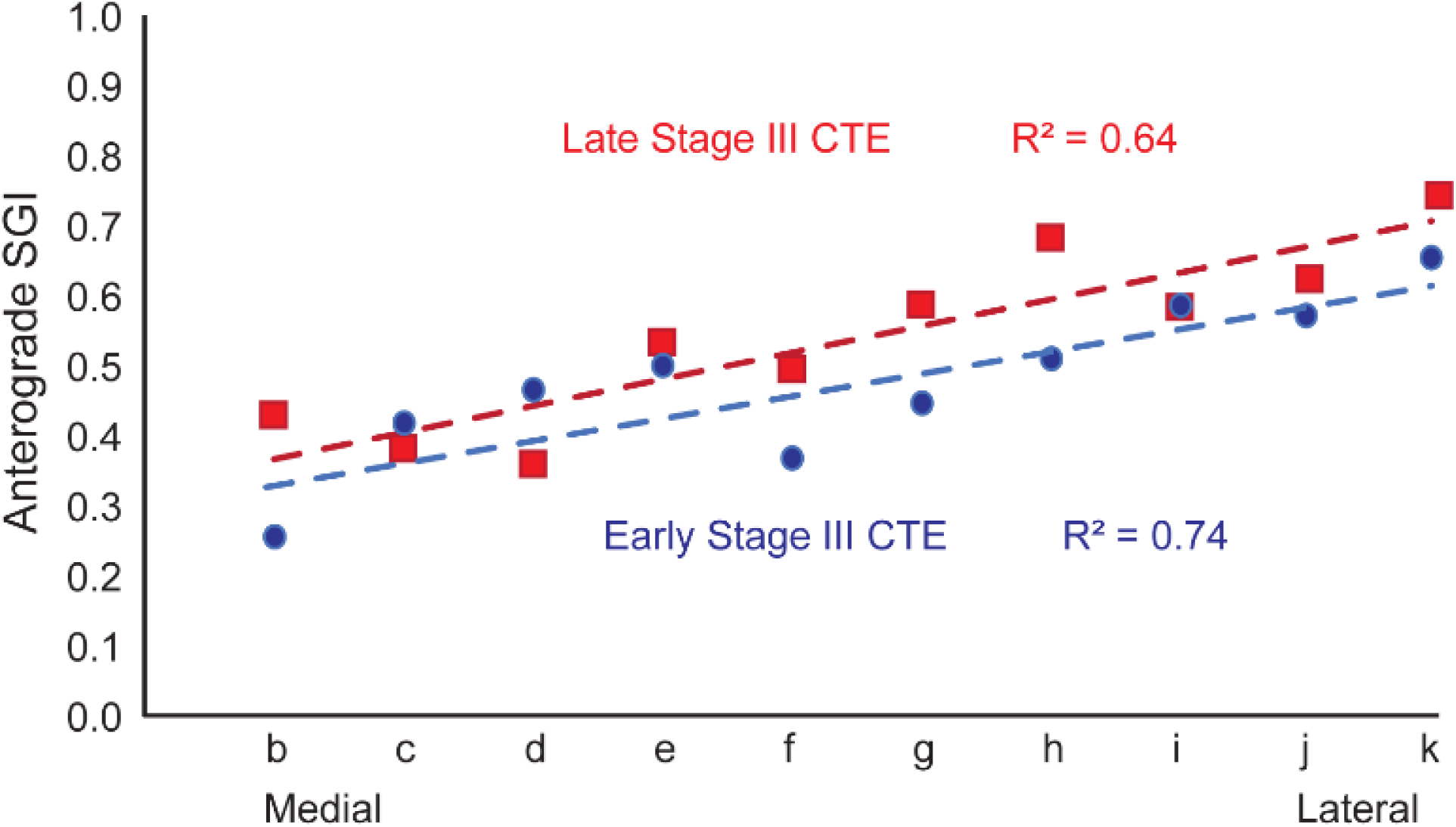
Laminar distribution of p-tau pathology in processes (excluding neuron bodies) as a proxy for anterograde pathways in stage III CTE. Early stage III CTE (blue circles); late stage III CTE (red squares).

The graded laminar pattern of p-tau pathology in axons in the anterograde direction appeared to be more subtle than the neurons labeled with p-tau pathology or connection patterns in animals, likely due to removal of degenerating axons by glia, a process that starts early after axon damage and has a variable duration ^25^; the timing of this process is not known for CTE, or the time interval between repetitive head impact and death. Similar trends in the spread of anterograde p-tau pathology were seen in the other cases (Supplementary Figs. 1 and 2).

### Comparison with findings in macaque studies

For comparison with connections in non-human primates we included findings of interconnections within the homologous MTL region of rhesus monkeys. The data were computed from cases with the use of gold-standard neural tracers injected in different parts of MTL and adjacent inferior temporal visual and superior temporal auditory association areas. Figure 4a shows a cross section with connections in MTL after injection of a bidirectional neural tracer in MTL A36 in a rhesus monkey. As seen in the magnified panels below, the predominant termination in A28 (4c) is in the middle-deep layers, reflecting a connection from an area with more elaborate laminar structure than the destination in A28. Fig. 4d shows distribution of terminations throughout the layers at a site close to the seed area, reflecting a lateral pattern of connections, while panels e and f show predominant termination in the upper layers, consistent with a pathway from an area with less elaborate to an area with more elaborate structure (feedback). The bidirectional tracer labels neurons as well, which are mingled in a heavy background of anterograde label (white label), are mapped in Fig. 4g. Projection neurons directed to area 36 are seen in the deep layers of A28 and 35 which have fewer layers, through a shift to both the upper and lower layers in an adjacent site of A36 (lateral pattern), and seen mostly in the upper layers in eulaminate visual area TE1 (red dots), as predicted by the Structural Model.

**Fig. 4:**
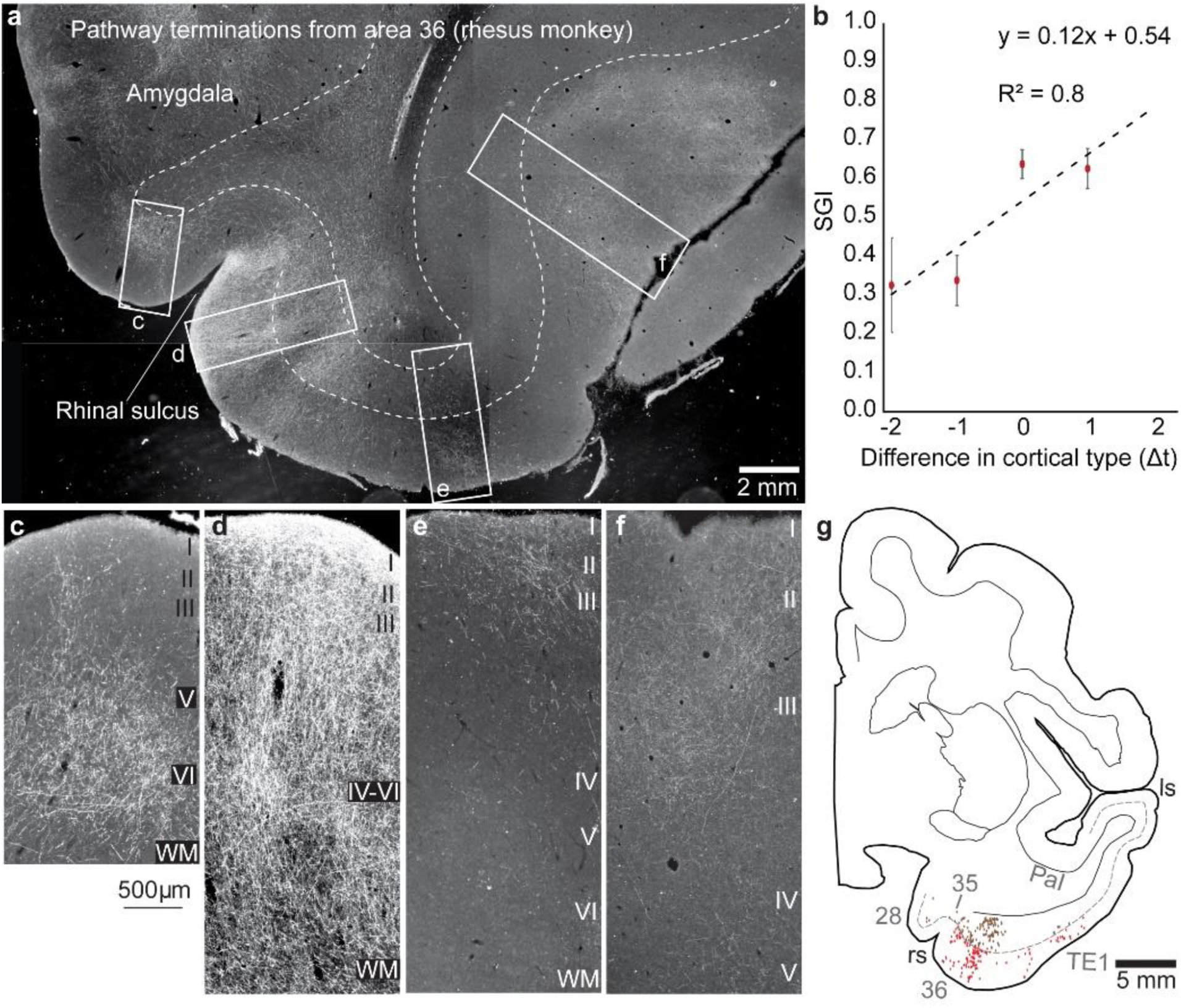
Interconnections of MTL areas in rhesus monkeys. **a**, Overview of representative coronal section through the temporal lobe at the level of the amygdala shows pathway terminations and projection neurons (white label) connected with medial temporal A36 (neural tracer was biotinylated dextran amine neural tracer (BDA)) injected in area 36. Magnified boxes (from a) **c-f**, Medial A28 is connected with A36 via the deep layers, d, shows connections in all layers and progressively in the upper layers of more lateral MTL sites. **b,** Quantitative analysis shows monotonic increase of projection neurons in the upper layers (SGI) from medial to lateral parts of MTL. Connection data were combined from 5 cases and expressed as a function of the difference in laminar type with respect to the injection site in each case. **g,** Projection neurons in the supragranular layers (red dots) and infragranular layers (brown dots) mapped in a cross section through MTL show a shift of projection neurons from the deep layers in A28, to both superficial and deep layers in adjacent MTL A36, and mostly in the upper layers of visual association area TE. The dotted line shows the upper part of layer V. ls, lateral sulcus; rs, rhinal sulcus. (a-f, darkfield photomicrographs)

Figure 4b combines connection data from 5 cases. Data were expressed as a function of the difference in laminar type with respect to the injection site in each case. The relational model makes it possible to combine data from different cases with injection of tracers (seed sites) in diverse temporal areas and to express the findings by quantitative analysis of labeled neurons in each area based on the structural relationship for pairs of linked areas in each case. Accordingly, on the x axis, 0 represents projection neurons from areas that are at about the same level as the seed site with respect to laminar structure, as in a lateral pattern (e.g., ^26^). Sites in increasingly lateral sites within MTL have progressively higher laminar differentiation than the site of origin (seed site), broadly called feedforward, with the x axis showing the range of values seen in increasingly lateral sites (positive numbers). Negative values on the x axis in Fig. 4b show the SGI from areas that have a less differentiated laminar structure than the injection site (seed area).

The graded patterns are not dependent on the distance between connected areas, as discussed in previous studies ^26^, and our observations here that p-tau pathology in the dysgranular insula, which is situated at a considerable distance from the seed site of A28, revealed a similar bilaminar pattern as the MTL site next to A28, which is also dysgranular (Fig. 1c; and Supplementary Fig. 4). In summary, the human CTE pathology in neurons and axons mimics the laminar patterns and density of cortico-cortical connections in monkeys.

### Propagation model

To illustrate how the Structural Model can account for the spread of p-tau pathology in CTE, we developed a propagation model to simulate stage III CTE and beyond. Figure 5 shows the relative distribution of p-tau pathology in supragranular layers (L2/3) and in the infragranular layers (L5/6) for the progression of the disease. We included five types of cortices that differ by degree of lamination. The first affected area is the cortex with the simplest laminar structure (agranular, type 1, akin to A28). All model cortices are connected to each other, with the strength of connection determined by the degree of similarity in laminar structure, as in cortico-cortical connections. The difference in degree of lamination also determines the laminar pattern of connectivity: the greater the difference in laminar structure for pairs of linked cortical areas the more asymmetrical the connectivity pattern, reflected in laminar distribution and density of connections. The more similar two connected areas are, the more symmetrical is the connectivity pattern. The difference in laminar architecture thus determines the extent to which the connectivity reflects feedforward/feedback versus lateral connections. In Figure 5, moving from left to right, the tauopathy progresses from an early to a late stage. In the first three stages, the supragranular layers are affected most (subplots a-1 to a-3), leading to SGI that increases with degree of lamination (subplots b-1 to b-3). The strength of connectivity decreases with degree of lamination (subplots c-1 to c-3). When the late stage is reached, more and more cortical areas serve as seeds for spreading p-tau, resulting in equalizing the SGI pattern (subplot b-4) and the density pattern (subplot c-4), as seen in CTE stage IV (Supplementary Fig. 5). Details of the model for propagation of p-tau pathology are in Methods.

**Fig. 5:**
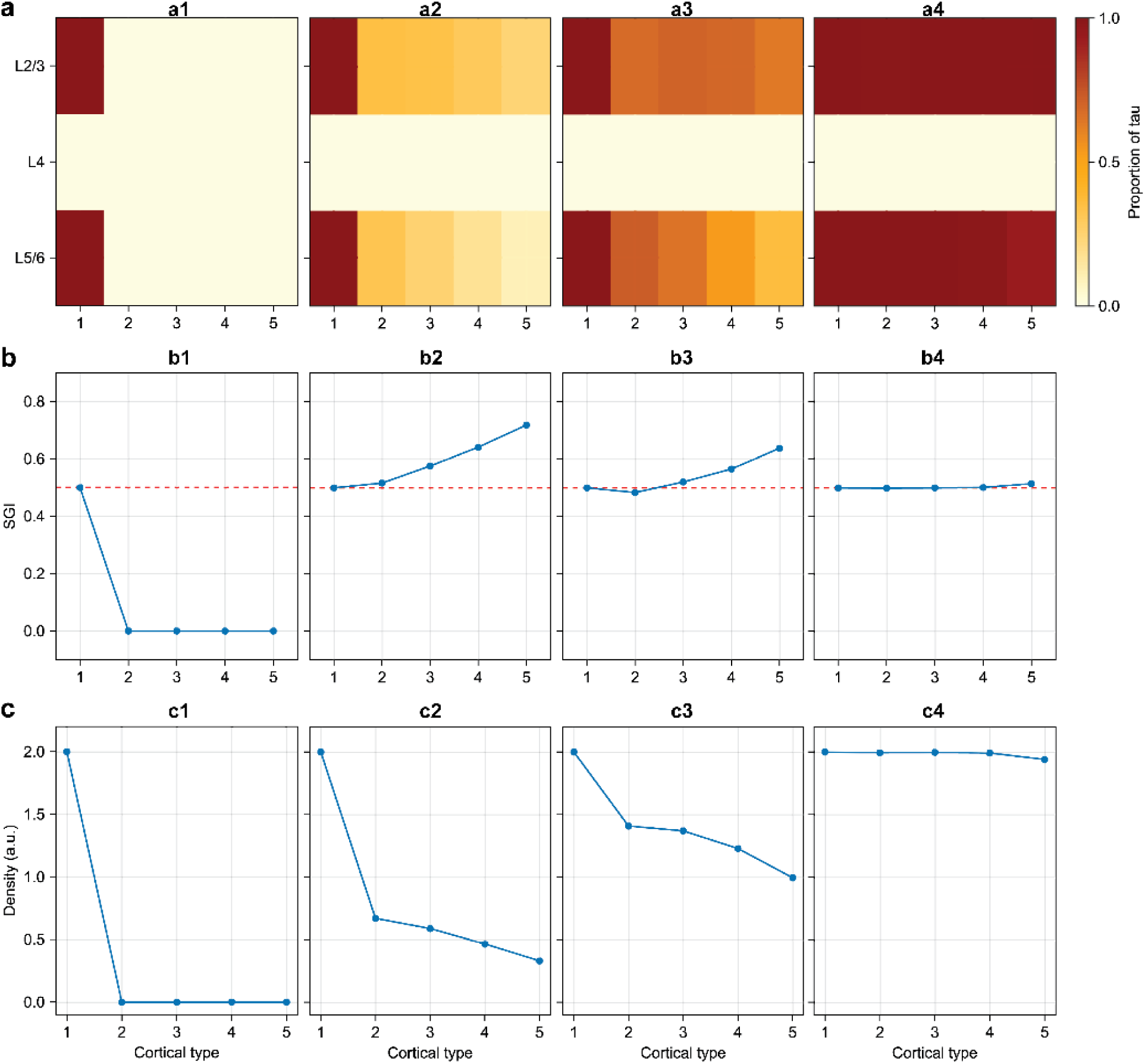
Simulation of propagation of p-tau across cortical types. Five cortical types were simulated, with the most limbic cortex (Type 1, agranular) being the seed area from which p-tau initially spreads. The disease advanced in stages, moving from left to right. Subplots a depict the relative uptake of p-tau in supragranular layers (L2/3) and infragranular layers (L5/6). Uptake is quantified as a proportion varying between 0 and 1: **a,** value of 1 means that all the neurons in the corresponding layer have p-tau. **b**, Subplots depict the corresponding supragranular gray index (SGI), which captures the extent to which the supragranular layers are preferentially stained with p-tau. Red dashed lines indicate SGI of 0.5, which corresponds to equal staining in supragranular and infragranular layers. **c**, Subplots depict the total amount of p-tau in the entire cortical area, which takes a maximum value of 2 when both supragranular and infragranular layers are fully stained (a.u.: arbitrary units). As the disease progresses, p-tau expression becomes increasingly uniform across L2/3 and L5/6, and also across cortices (subplots a-4, b-4, and b-4).

### Differential pattern of label in the depths of sulci in early CTE (stages I, II): columnar connections

The pattern of cortical p-tau pathology in MTL in CTE stage III follows cortico-cortical circuit rules, in which the only layer that does not include labeled neurons is layer 4 (Figs. 1 and 2; and Supplementary Figs. 1, 2). By contrast, within the focal label in the depths of sulci in early CTE (stages I and II), ∼10% of p-tau labeled neurons were found in layer 4, and included labeled neurons in the other cellular layers as well (Fig. 6). This pattern resembles a different type of circuitry, namely connections within a cortical column, which includes participation of layer 4 neurons (reviewed in ^27,28^). Layer 4 neurons have short axons that do not leave the cortical mantle, as noted first by Ramon-y-Cajal ^29^, but participate in connections within a column and connections across the adjacent 1-2 columns. In both types of circuits the termination of axons involves all layers.

**Fig. 6:**
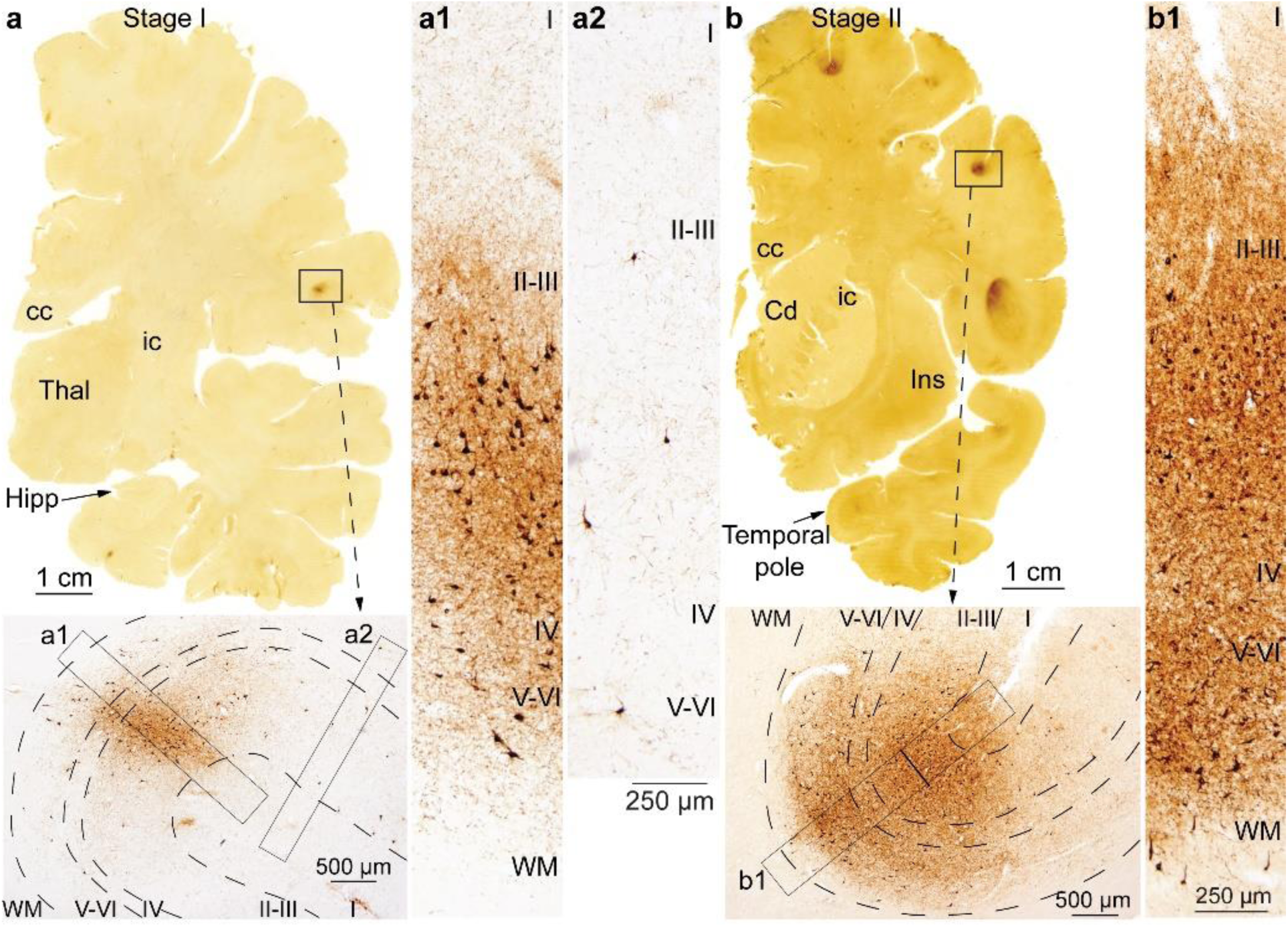
Expression of p-tau pathology in dorsal and lateral cortices in early CTE stages. **a**, Cross section through the temporal lobe at the level of the anterior hippocampus immunostained against p-tau in stage I CTE. Outlined region shows initial area of injury in the depths of a sulcus, magnified below to highlight labeled neurons in all layers, including layer 4. **a1, a2**, Outlined regions are shown at higher magnification on the right side of panel a. In the depths of the sulcus in a1 there are labeled neurons in all layers, a pattern that resembles a different type of circuitry, namely connections within a cortical column. Columns in nearby areas (a2) do not include neurons in layer 4, a pattern that resembles cortico-cortical connections. **b,** Overview of cross section through the temporal lobe at the level of the temporal pole immunostained against hyperphosphorylated tau in stage II CTE. Outlined region (box) shows early p-tau pathology at higher magnification shows labeled neurons in all layers, including layer 4. **b1**, Outlined region is shown at higher magnification on the right side of panel b.

## Discussion

Our findings revealed a striking parallel in the pattern of p-tau pathology in human MTL in CTE and cortico-cortical connections in monkeys. Significantly, the p-tau pathology pattern conforms to the predictions of a theoretical model that uniquely reveals the direction of connections, and in this instance p-tau pathology and its progression in successive MTL sites by laminar distribution of affected neurons and axons. The fundamental predictions of the model were supported, as previously seen in the connections in diverse animal species and systems (reviewed in ^1,3^).

Another prediction was the absence of neurons with p-tau pathology in layer 4, which do not participate in cortico-cortical connections but interconnect neurons up and down their particular narrow column ^27,28^, as seen in the earliest stages of CTE in sporadic sites within the depths of sulci. These predictions were consistently substantiated for the first time in the human cortex through quantitative analyses in single cases and sections attesting to the robustness of the findings.

According to the reciprocity rule of cortico-cortical connections, the primate A28 projects to several MTL areas and receives projections from these areas in non-human primates ^22^ (Fig. 4). P-tau pathology was also seen in neurons and axons, consistent with reports that misfolded tau complexes are internalized in dendrites and axons and travel in both directions (e.g., ^30^). Accordingly, neurons with p-tau pathology in CTE stage III showed graded differences in laminar distribution and density in MTL. In areas that abut A28, neurons with p-tau pathology were densely distributed in more layers (layers 2-3 and 5-6), reminiscent of lateral connections in monkeys (see ^26^). In sequential lateral MTL areas, sparser neurons with p-tau pathology were found progressively in the upper layers, as seen with feedforward projection patterns in animal connectional studies, and consistent with model predictions.

The medial part of A28 showed the earliest and densest p-tau pathology in CTE stage III, operationally establishing it as the cortical seed site. In the example in Figure 2 of early stage III CTE, p-tau pathology was overwhelmingly seen in the deep layers. From A28, a graded pattern of p-tau pathology could be traced in the anterograde direction. By circuit rules, the deep layers of A28 project to more lateral parts of MTL via feedback-like pathways that target all layers of the nearby dysgranular sites, and progressively the upper layers in more lateral areas that have increasingly differentiated laminar structure. By circuit rules and application of a computational model here, these findings provide concrete evidence of the direction of p-tau pathology in time, with the implication that pathology affects all areas connected with A28 (the seed site) in a graded manner. In time, as areas close to the seed site become increasingly affected with p-tau pathology, they turn into seed areas themselves, affecting all areas they are connected with, in a chain reaction leading to the inexorable progression of the disease with widespread distribution of pathology in cortical layers by CTE stage IV (e.g., ^17^, Supplementary Fig. 5), and as revealed by the computational progression model here.

Progression of pathology via connections has been suspected in animal models of traumatic brain injury (e.g., ^31^) and in neurodegenerative diseases (e.g., ^13,32–35^) such as Alzheimer’s disease (AD) ^14,15^. In the latter, the seed cortical area is the ‘transentorhinal’ region with pathology seen more medially in progressive stages of Alzheimer’s disease ^19^, with the hippocampus affected after the cortical entorhinal region. By contrast, in CTE III the seed cortical area is the medial part of A28, with progression of p-tau pathology to more lateral sites within MTL, revealing that in AD and CTE, p-tau pathology proceeds in opposite directions.

Hypotheses have been advanced that pathology in neurodegenerative diseases may spread by a non-circuit basis through microglia or exosomes ^36–38^. Our findings cannot rule out some contribution by such mechanisms. However, the ordered laminar distribution of p-tau pathology in both neurons and axon terminals strongly points to circuit mechanisms as the primary mode of spread of the pathology, since any substantial non-circuit spread would have obscured the consistent laminar distribution of p-tau pathology found here.

Previous studies have also distinguished the early focal, perivascular p-tau pathology in the depths of sulci in frontal or temporal lobes in CTE stages I - II from the broad p-tau pathology that occurs in the MTL in CTE stage III (e.g., ^17,39^). The early focal perivascular p-tau pathology likely reflects the areas of greatest physical deformation during head impact injury (reviewed in ^35^), whereas the progressive diffuse pathology found in the MTL in stage III likely represents a spreading phenomenon ^40^. A key distinction noted for the first time here, is the presence of neurons with p-tau pathology in layer 4 in the depths of sulci in early CTE, consistent with the distinct circuitry within cortical columns (reviewed in ^27,28^) but not cortico-cortical connections.

### Functional implications

The systematic variation of the cortex can be traced to development and has implications for cortical evolution ^7,41^. The linkage of cortico-cortical connections to the systematic variation of the cortex has provided a set of rules to predict connections and probe direction and progression of neurodegenerative diseases, as applied here for CTE. The Structural Model also predicts the cortical plasticity-stability continuum ^42^ and likely vulnerability to disease ^1,7^. By this measure, the agranular (limbic) A28 and adjoining dysgranular areas are the most plastic areas in MTL, consistent with their involvement in learning, memory and emotions. The plasticity of limbic areas also renders them vulnerable to disease, as seen here for CTE.

The involvement of A28 in the systematic p-tau pathology in CTE has important implications for function. A28 receives pathways from high-order sensory and other association cortices ^43^, projects to hippocampus, receives the hippocampal output in its deep layers, and projects to high-order association cortices (e.g.,^44^). These pathways endow A28 with information to process memories in context ^45,46^, engage in complex spatial tasks, and form relational associations for abstract processes within a cognitive and mnemonic domain ^47–49^. The pathology in MTL likely disrupts pathways from hippocampus and other cortical areas. One pathway that originates in medial prefrontal A25 innervates heavily the deep layers of A28, is associated with autobiographical memory and emotions ^50^. These processes are severely affected in psychiatric diseases, including PTSD and depression, which are also experienced by young people later diagnosed with CTE ^51^. The strong parallels in p-tau pathology with cortico-cortical connections points to the potential that repetitive head impact may damage early diverse cortical pathways that converge on A28 and collectively contribute to the systematic p-tau pathology seen in the MTL in CTE stage III.

## Figures and Figure Legends

**Supplementary Fig. 1:**
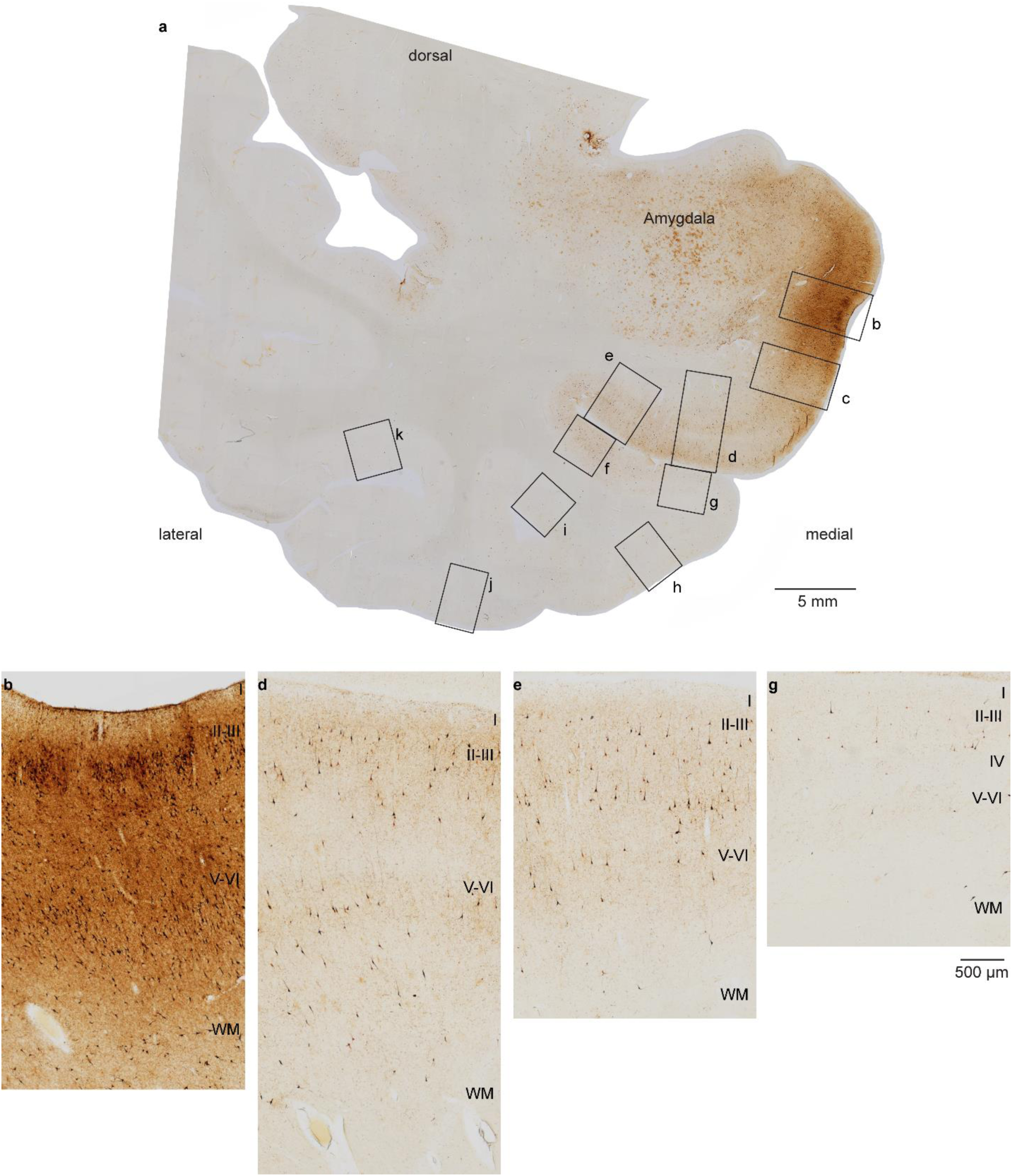
P-tau pathology in temporal cortices in late stage III CTE. **a**, Overview of representative coronal section through the medial temporal lobe of case K-281 at the level of the amygdala immunostained against p-tau. **b-g,** High magnification of regions (outlined in a) highlight the graded pattern of the laminar distribution of neurons with p-tau pathology that parallels patterns of laminar connectivity between medial A28 (b) through adjacent dysgranular and progressively eulaminate areas with more elaborate laminar structure.

**Supplementary Fig. 2:**
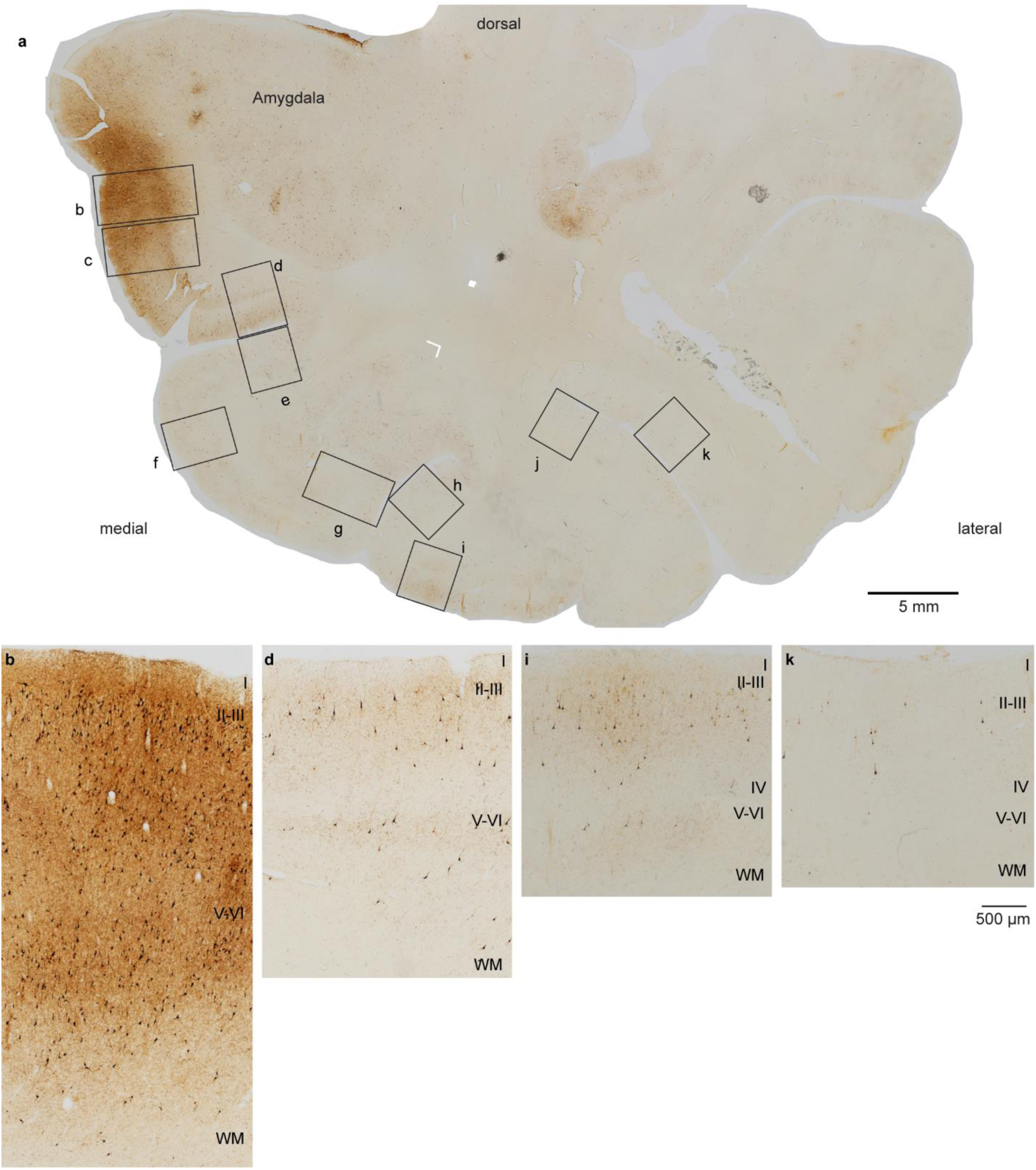
P-tau pathology in temporal cortices in late stage III CTE. **a**, Overview of coronal section through the medial temporal lobe (case K-372) at the level of the amygdala immunostained against hyperphosphorylated tau. **b-k**, High magnification of regions outlined in a (b-k) highlight the graded pattern of the laminar distribution of neurons with p-tau pathology show the same pattern as in Figs. 1-2; p-tau pathology here parallels patterns of laminar connectivity between agranular/dysgranular medial and progressively more eulaminate lateral temporal areas.

**Supplementary Fig. 3:**
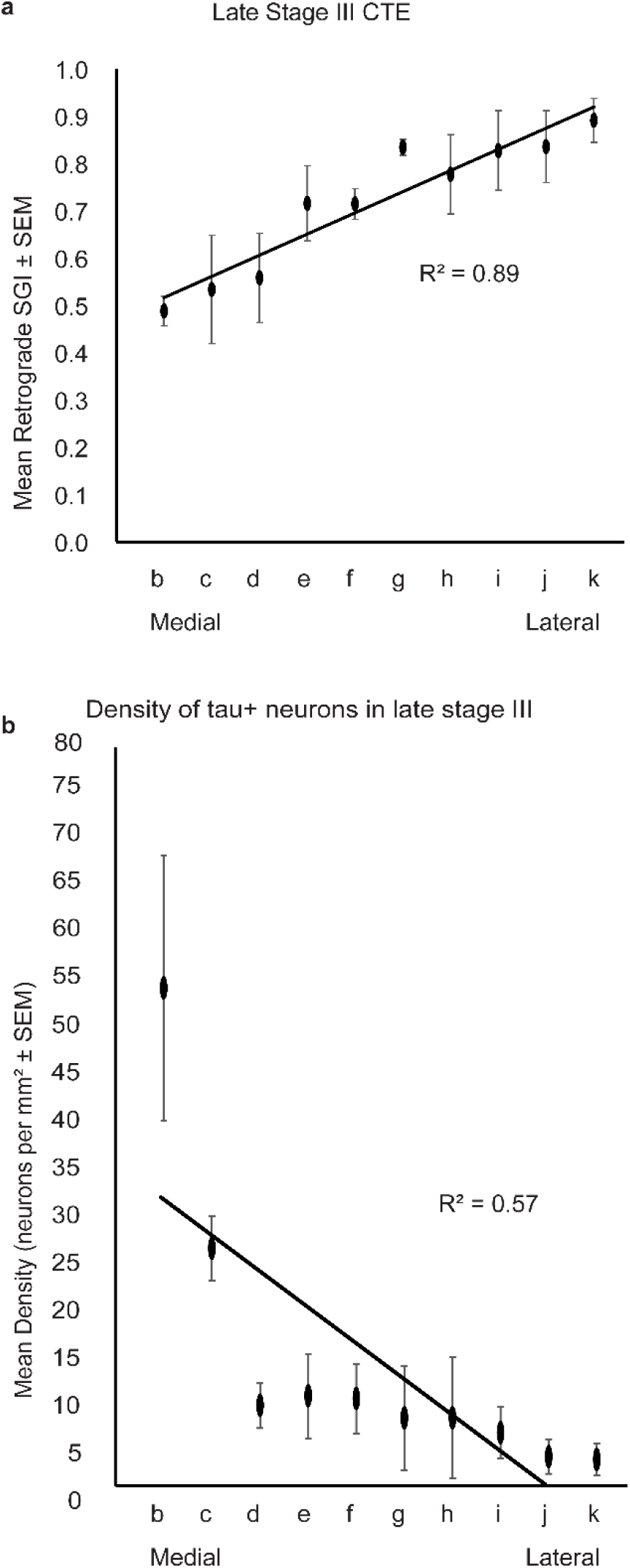
Laminar distribution and density of p-tau pathology in temporal cortices in late stage III CTE. **a**, Quantitative analysis shows monotonic increase of neurons with p-tau pathology in the upper layers (SGI) from a medial to lateral parts of MTL, averaged for three cases with late stage III CTE. **b**, Quantitative analysis of the density of neurons with p-tau pathology in the supragranular layers, averaged from three cases with late stage III CTE, shows gradual decrease from A28 to successive adjacent lateral sites within MTL, which have progressively higher differentiation of layers.

**Supplementary Fig. 4:**
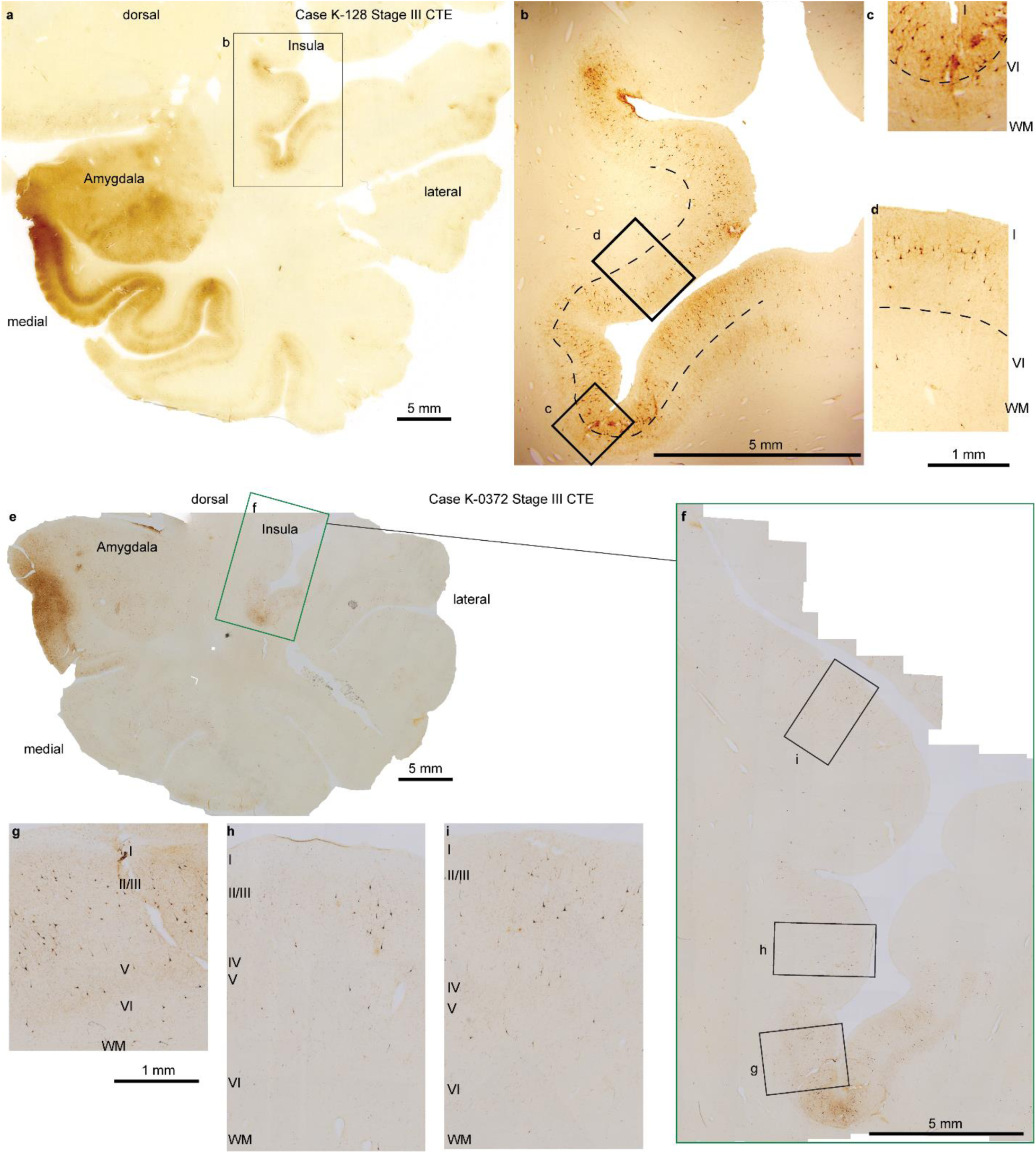
P-tau pathology in insular cortices in late stage III CTE. **a**, Overview of cross section through the medial temporal lobe at the level of the amygdala immunostained against hyperphosphorylated tau (case K-128). **b,** High magnification of region outlined in a (b), shows similar multi-laminar distribution of p-tau in the insula as at medial temporal sites situated close to the seed side of the entorhinal cortex, even though the two sites are at a considerable distance from each other. **c-d**, High magnification of regions in b (from a) show the graded pattern of the laminar distribution of neurons with p-tau pathology in the insula, which parallels patterns observed in MTL. **e-i**, Progressively higher magnifications of hyperphosphorylated tau expression in insular cortices of case K-0372 show parallel patterns of p-tau pathology observed in MTL. The dotted line in b-d shows the upper part of layer V.

**Supplementary Fig. 5:**
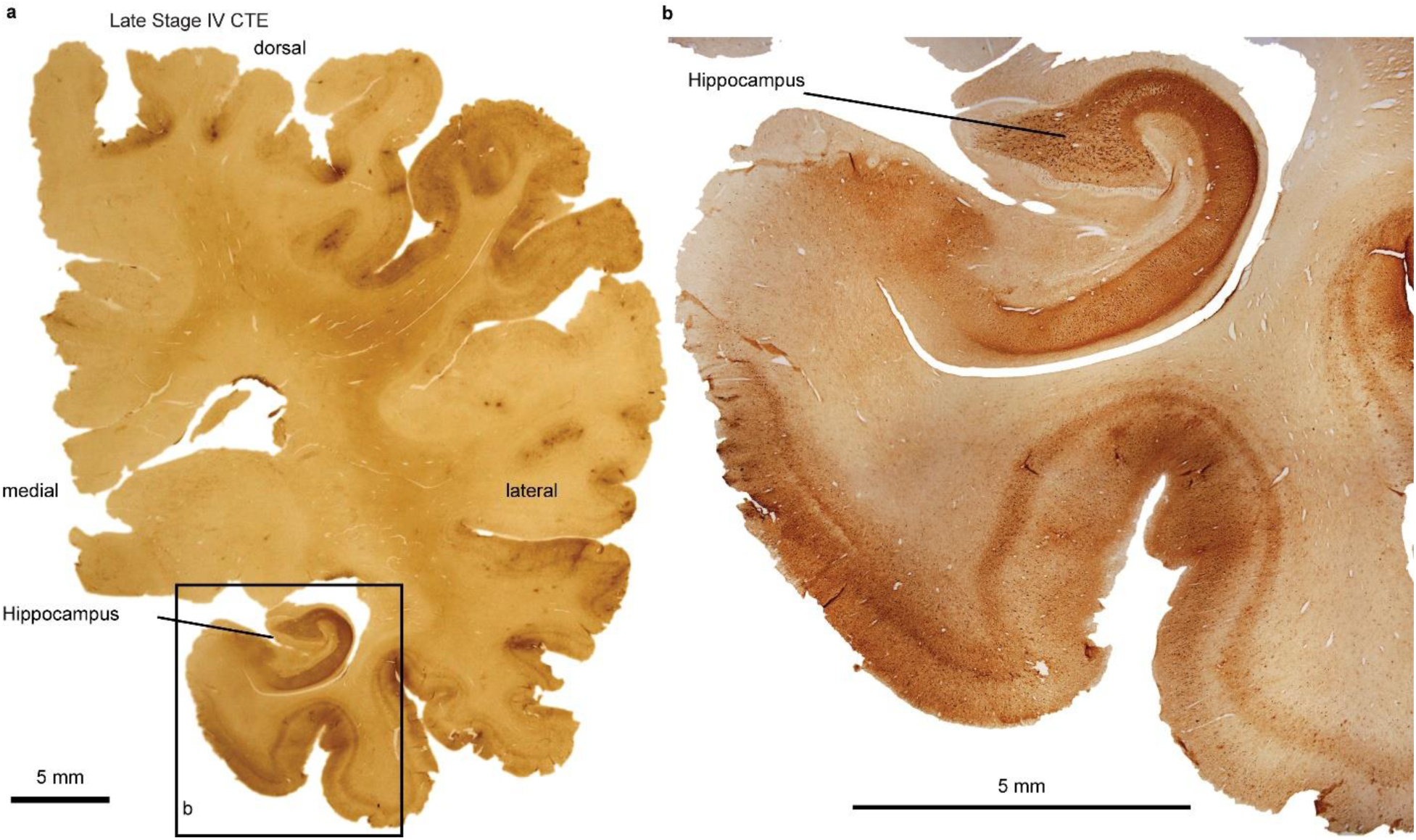
Widespread expression of p-tau pathology in the cortex in CTE stage IV. **a**, Overview of cross section through the medial temporal lobe at the level of the hippocampus immunostained against hyperphosphorylated tau. **b**, High magnification of region outlined in box b (from a) shows extensive spread of neurons with p-tau pathology in MTL and throughout the cortex in a late stage of the disease.

## Materials and Methods

### Human postmortem brain and tissue preparation

The study was approved by the Institutional Review Board of Boston University. Data of human *postmortem* brain tissue are summarized in Table 1. CTE cases were from the BU CTE Brain Bank (N=12; Supplementary Table 1). Human tissue from the CTE Brain Bank was fixed in periodate–lysine–paraformaldehyde (PLP) and stored at 4 °C until block preparation and sectioning. In addition, we examined the cytoarchitecture of temporal cortices in *postmortem* human brain tissue from neurotypical individuals (N = 3, one female) to determine the laminar structure and cortical types of temporal areas, using Nissl histological staining, as described in detail earlier ^21,41,52^. Donated *postmortem* human brains from the National Disease Research Interchange (NDRI) were immerse-fixed in 10% formalin. We then sliced each cerebral hemisphere in coronal slabs of 1 cm thickness, photographed the anterior and posterior surfaces of each slab and post-fixed them in 10% formalin for 2–4 days. After post-fixation, we matched the slabs based on atlases of the human brain ^53,54^. We cryoprotected tissue slabs in a series of sucrose solutions (10–30% in 0.01 M PBS), and temporal blocks were frozen in −75°C isopentane (Thermo Fisher Scientific, Pittsburg, PA, United States) for rapid and uniform freezing ^55^. We then cut blocks on a freezing microtome in the coronal plane at 50 μm and 10 consecutive series of sections were collected. Sections were stored in antifreeze solution (30% ethylene glycol, 30% glycerol, 40% PB 0.05 M at pH 7.4 with 0.05% azide) at −20°C for future assays.

**Supplementary Table 1.**
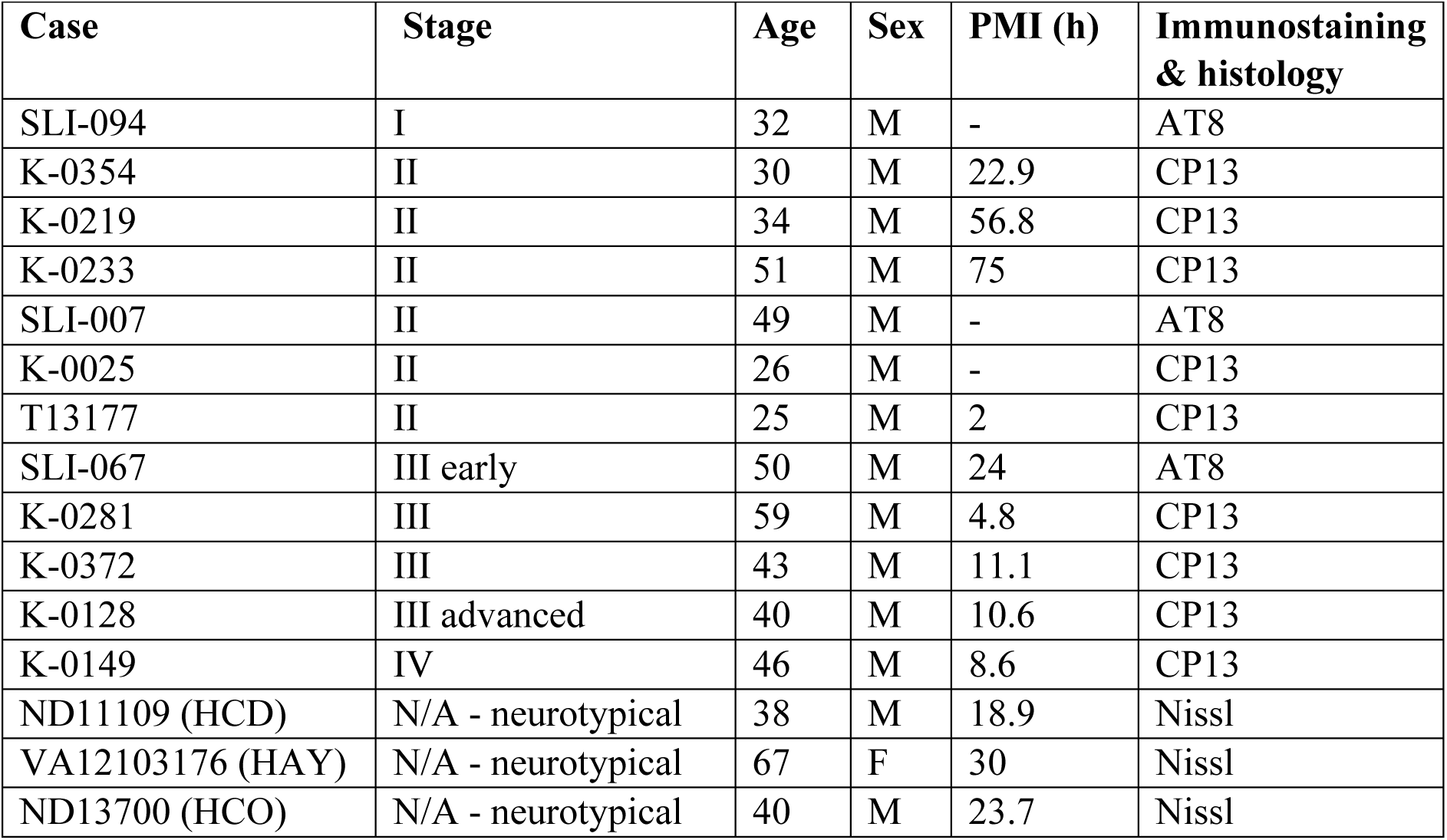
*Postmortem* human brain tissue information.

### p-Tau visualization in human postmortem tissue sections

For AT8 and CP13 phospho-tau histopathology, tissue blocks were embedded in paraffin for sectioning. Serial 10 μm sections were cut and mounted for immunohistochemistry, as described ^56–58^. Briefly, mounted sections were incubated overnight at 4 °C in primary antibody AT8 (a mouse monoclonal antibody directed against phosphoserine 202 and phosphothreonine 205 of PHF-tau; Pierce Endogen, Rockford, IL; 1:2000), or CP13 (a monoclonal antibody directed against phosphoserine 202 of tau, considered to be the initial site of tau phosphorylation in neurofibrillary tangle formation; 1:200; courtesy of Peter Davies) followed by biotinylated anti mouse secondary antibody. P-tau was visualized with a 3-amino-9-ethylcarbazol HRP substrate kit (Vector Laboratories, Burlingame, CA). Sections were coverslipped with Permount medium (Thermo Scientific, Rockford, IL).

### Neuropathological evaluation and stage classification of CTE

Diagnosis and stage classification were based on characterization of pathognomonic lesions, as described in detail in published criteria for the neuropathological diagnosis of CTE ^59, 60^. CTE is characterized by progressive buildup of hyperphosphorylated tau (p-tau) in neurofibrillary tangles and neurites that initially surrounds small blood vessels at the sulcal depths of the cerebral cortex and later is seen in different cortical and subcortical areas. The density and regional deposition of p-tau determines four pathological stages of CTE, ranging from stage I (mild) to stage IV (severe), with gradually increasing and more widespread p-tau deposition, with fibrils that have distinctive molecular structural configuration unlike the configurations observed in aging, Alzheimer’s disease, or other p-tau pathology ^18^.

### Animals, surgery, tracer injections, and tissue preparation

We used available *postmortem* rhesus monkey (*Macaca mulatta,* ∼2-3 years, n = 3, female = 1) brain tissue for architectonic and pathway analyses (Supplementary Table 2). Animal protocols were approved by the Institutional Animal Care and Use Committee at Boston University School of Medicine and Harvard Medical School, in accordance with the ILAR *Guide for the Care and Use of Laboratory Animals* (publication 80–22 revised, 1996). We designed experimental procedures to minimize animal suffering and to reduce the number of animals needed for research, by placing multiple distinct tracers in each case, which we used for this and other unrelated studies.

Tracer injection sites and quantification of projection neurons have been described in detail in previous studies and will be briefly described here ^23^. First, we obtained high-resolution magnetic resonance imaging (MRI) scans of the brain *in vivo* for surgical planning of tracer injections. Animals were sedated with ketamine hydrochloride (10-15 mg/kg, i.m.), then anesthetized with propofol (loading dose 2.5-5 mg/kg, i.v.; continuous infusion rate 0.25-0.4 mg/kg/min), and placed in a stereotaxic apparatus (1430M; David Kopf Instruments) for the scan. We used MRI scans to calculate stereotaxic coordinates using the interaural line as reference for five injections of neural tracers in several temporal cortical areas (Table 2).

We performed surgery for tracer injections under general anesthesia (isoflurane, to a surgical level) with continuous monitoring of vital signs. We injected one or three tracers in each monkey, as shown in Supplementary Table 2. The dyes injected included fluorescent and other tracers: Fast Blue (FB, case AT, 3% dilution, 2 µl; Polysciences), Diamidino Yellow (DY, case AT, 3% dilution, 4 µl,), Fluoro-ruby (FR, case BC, (10% solution, mixture of 10 kDA and 3 kDA, 4 µl; Invitrogen), and Biotinylated Dextran Amine (BDA, cases AV, AT: equal parts 10% 10 kDA and 10% 3 kDA, 5 µl; Invitrogen). Dextran amines with molecular weight 10 kDA (BDA in AT and AV) are optimal for anterograde labeling of axonal projections and their terminations, while the 3 kDA variant is optimal for retrograde labeling of cell bodies and proximal dendrites ^61–63^. Post-operatively we monitored animals and gave antibiotics and analgesics.

After a survival period of 18-20 days, the animals were anesthetized with a lethal dose of sodium pentobarbital (∼50 mg/kg, i.v., to effect), and perfused transcardially with 4% paraformaldehyde in cacodylate buffer or in 0.1M PB at pH 7.4 (cases AT, AV), or 4% paraformaldehyde and 0.2% glutaraldehyde in 0.1M PB, pH 7.4 (case BC). The brains were removed, photographed, cryoprotected in ascending concentrations of sucrose solutions (10-25% sucrose in 0.1M PB, pH 7.4, with 0.05% sodium azide; Sigma Aldrich), and frozen in -80°C isopentane ^55^. We used a freezing microtome (AO Scientific Instruments/Reichert Technologies) to section brains in 50 µm-thick coronal sections, for case BC, or 40 µm for cases AT, AV, producing ten matched series of sections that were stored in antifreeze (30% ethylene glycol, 30% glycerol, 0.05% sodium azide in 0.05M phosphate buffer, pH 7.4), until further processing.

**Supplementary Table 2.** List of rhesus macaque cases and analyses performed.

| Case | Sex | Age (years) | Hemisphere | Injection site | Tracer | Analysis of tracer label |
| --- | --- | --- | --- | --- | --- | --- |
| AT | Female | 2 | Right | A36 | BDA | Retrograde/anterograde |
| AT | Female | 2 | Right | TE1 | FB | Retrograde |
| AT | Female | 2 | Right | TPro | DY | Retrograde |
| AV | - | - | Left | TE1/Ts1 | BDA | Retrograde/anterograde |
| BC | Male | - | Right | STG | FR | Retrograde |

### Tracer visualization in rhesus monkey tissue sections

In the cases with injection of fluorescent tracers, sections were mounted immediately after cutting, dried, coverslipped with Krystalon (Millipore), and labeled neurons were mapped directly, as described ^64^. Briefly, sections used to visualize neurons retrogradely labeled for tracers (FB, DY or FR) were viewed under a microscope with epifluorescence attachment (Nikon Optiphot) that includes an analog-digital microscope/computer interface (Nikon, Optiphot/Austin), for mapping and counting labeled neurons and axon terminals with custom-made software or using a commercial system (Neurolucida, v.10.40 Microbrightfield).

To visualize projection neurons labeled by BDA we incubated free-floating sections for 1 h at 4°C in 0.05M glycine (Sigma-Aldrich) and preblocked for 1 h at 4°C in 5% normal goat serum (NGS; Vector Laboratories), 5% bovine serum albumin (BSA; Sigma-Aldrich), and 0.2% Triton-X (Sigma-Aldrich) in 0.01M PBS. Sections were then incubated for 1 h at 25°C with avidin-biotin horseradish peroxidase (AB-HRP; Vectastain Elite ABC kit, Vector) at a 1:100 dilution in PBS, followed by three rinses in PBS (10 min each), and tracer was visualized with incubation for 1–3 min in diaminobenzidine (DAB substrate kit, Vector). Sections were mounted on gelatin-coated glass slides, dried, counterstained with Nissl (thionin stain, every other section) as previously described ^65^, and coverslipped with Entellan (Sigma-Aldrich).

### Quantitative analysis of p-tau labeling in the human cortex in CTE

We quantified overall p-tau labeling density across layers and cortices in immunostained coronal sections from cases with early (I and II) and advanced (III) CTE stages. We estimated the laminar p-tau content from images of representative columns from each area in the human frontal and temporal cortices captured at the light microscope (Olympus BX 51), under brightfield illumination, with a CCD camera (Olympus DP70), connected to a workstation running imaging software (DP Controller, cellSens Standard 3.2, Olympus). We captured images using 4x, 10x, and 20x lenses with the same light exposure to minimize background variability, as described ^64^. We imported images into ImageJ (v2), for further analyses of the overall signal density, the number of labeled neurons, as a proxy for retrograde labeling, and the density of p-tau signal excluding neurons, as a proxy for anterograde labeling, since abnormal tau travels primarily through neurons and labels axons ^11,56,66,67^.

#### Overall and laminar signal density

To measure signal density, we converted images into 8-bit gray scale, and inverted them, so that signal intensity measurements ranged from 0-255 ^64^. We measured levels of background staining in gray matter regions with no specific immunostaining within each image and subtracted background pixel values from each image to eliminate staining inconsistencies, due to experimental variability. We obtained optical density (mean gray value) and surface area measurements from each layer within frontal and temporal cortical gray matter columns and used these to additionally estimate the integrated density of the signal (mean gray value / area). We additionally used image thresholding to estimate the area fraction of p-tau labeling for each image, as an independent measure of signal density.

#### Laminar distribution and density of labeled neurons (retrograde)

We computed the number of labeled neurons across layers in representative cortical columns within frontal (for early CTE stages I and II) and temporal (for CTE stage III) cortices. To describe the laminar distribution of labeled neurons within and across cases, we expressed laminar specificity as the percent of labeled neurons originating in the upper layers (II-III), referred to as the supragranular index (SGI) for each site.

#### Laminar distribution and density of p-tau immunolabeling in axons and terminals (anterograde)

We estimated signal density excluding the signal from labeled neuron bodies, as a proxy for axonal p-tau signal. We segmented images using ImageJ to produce a binary image mask that facilitated extraction of only the pixels within cell bodies of interest and exclusion of other aspects of the image, as described ^68^. The resulting mask contained an outline of all p-tau+ cells. Each manually segmented binary mask was then used to extract image pixels from the matched grayscale original image using MATLAB (2019b, MathWorks, Natick, MA). To perform the optical density analysis, we normalized each extracted image so that the maximum brightness was 1. For each image, the average p-tau intensity per pixel was computed for the neuropil and was used to estimate the density ratio of signal in the superficial, deep, and all layers to express the supragranular index (SGI) as labeling in superficial / all layers.

### Analysis of anterograde and retrograde labeling in the rhesus monkey cortex

We mapped cortical pathways directed to medial and lateral temporal areas in a series of coronal sections through the medial and adjacent inferior temporal (visual) and superior temporal (auditory) association and multimodal areas in the depths of the superior temporal sulcus, using exhaustive plotting in 1 out of 20 coronal sections through the ipsilateral hemisphere, as described ^23^. In each series of sections, we mapped all labeled cortical neurons in the temporal cortex in the superficial (II-III), and deep (V-VI) layers, at 200x with brightfield or epifluorescence microscopy (Olympus BX60) and a semi-automated commercial system (Neurolucida v.10.40, MicroBrightField). We normalized estimates by dividing the number of labeled neurons in a specific area by the total number of labeled neurons in the temporal lobe of each case, to account for likely variation across cases in the size of injection sites, tracer transport dynamics, and immunolabeling, even when identical procedures are used. We summed labeled neurons across sections for each cortical area in each case to describe the laminar distribution of labeled neurons within and across cases and expressed laminar specificity as the percent of labeled neurons originating in the upper layers, the SGI. We estimated the mean and standard error of the mean of these percentages across cases.

We studied anterograde tracer termination patterns qualitatively in cases AT and AV (tracer injections in A36 and TE1/TS1) by analyzing the efferent projections from the injection sites to other temporal areas using ImageJ. We captured darkfield images using an Olympus optical microscope (BX53) with a CCD camera (Olympus DP74) connected to a commercial imaging system (DP Controller, Olympus). In ImageJ, we used the auto threshold feature and the moments method to analyze the images, as this best represented the signal of the tracer. We used the binary images to obtain the measurement for area fraction, which is the area of the image that was occupied by the signal. We acquired the area fraction measurement for the column (width of 150µm and height differed among areas) with the highest intensity within each area. We then computed the proportion of area fraction of each area of the temporal lobe by dividing the measurement recorded per area over the total amount of area fraction measured in this case.

### Statistical analysis

We used linear regression analysis to fit a line through each set of observations to examine the relationship between p-tau labeling and changes in cortical laminar structure. The fit can best predict how the laminar distribution and density of p-tau labeling can be affected by the gradual changes in cortical laminar architecture. We report significance and F values for 95% confidence level.

### Propagation model

We modeled the propagation of p-tau using a simple bounded integration model. The increment of the proportion of p-tau p in a given layer m of cortex area A at time step t given by

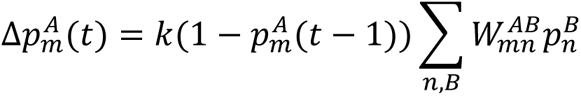

Where 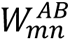 is the strength of a connection to layer *m* of cortical area *A* from the layer *n* in a connected cortical area *B*. The sum is taken over all connected cortical areas (*B*) and layers (*n*). For the simulation here, all cortices are assumed to be connected to each other. Note that local connectivity within a cortical area is omitted for simplicity. Rate constant *k* determines the speed of propagation.

We implemented the rules of the relational Structural Model by letting the difference in degree of lamination D regulate the laminar connection strength. The degree of lamination D was specified by a score varying from 0 to 1, with 0 corresponding to the most limbic cortex, and 1 corresponding to the most eulaminate cortex. The connection to an infragranular layer (i) in cortex A from a supragranular layer (s) in a connected cortex B is given by:

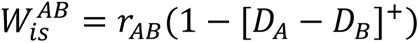

Similarly, the connection to a supragranular layer (s) in A from an infragranular layer (s) in B is given by:

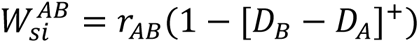

The expression [*x*]^+^ = *x* when *x* > 0 and is 0 otherwise. The term *r*_*AB*_ determines the maximum possible strength of the connection.

If cortex A is maximally limbic (*D*_*A*_ = 0) and cortex B is maximally eulaminate (*D*_*B*_ = 1) then 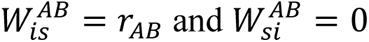, corresponding to a pure feedforward projection from B to A. For the reverse situation, when A is maximally eulaminate (*D*_*A*_ = 1) and cortex B is maximally limbic (*D*_*B*_ = 0) then 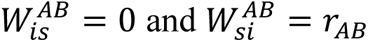, corresponding to a pure feedback projection from B to A. If the two cortical areas have the same degree of lamination, then both feedback and feedforward projections have equal and maximal strength (*r*_*AB*_), corresponding to a ‘lateral’ pattern that involves more layers. Intermediate levels of difference in degree of lamination produce biased patterns that are either more feedforward or more feedback, depending on the sign of the difference.

The maximal strength of a connection *r*_*AB*_ is not directly determined by the Structural Model. For many cortico-cortical connections, the strength covaries roughly with distance, but this does not always apply, given that some distant cortices can have strong connections. For simplicity we assumed *r*_*AB*_ is given by a Gaussian function of the difference in degree of lamination:

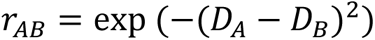

Simulation of the above model was run for 30 timesteps, with *k* = 0.1. The results for timesteps 1, 5, 10, and 30 are shown in Figure 6. Simulations were run using the Julia programming language.

## Acknowledgements

This work was supported the National Institutes of Health, National Institute of Mental Health grant no: R01MH117785 (HB), R01MH136013 (HB, BZ) and Neurological Institute of Disorders and Stroke (U01NS086659, U01NS093334, U54NS115266 (AmK)).

## Author contributions

Conceptualization: HB, MAGC, BZ; Experimental procedures: HB, MAGC, JB, AmK, BZ; Neuropathology: AmK; Formal analysis: HB, MAGC, JB, BZ; Computational modeling: YJ, Funding acquisition: HB, AmK; Writing – original draft: HB; Writing – review & editing: HB, MAGC, JB, AmK, YJ, BZ.

## Data availability

Data supporting the findings of this study are either included in the manuscript or are available from the corresponding author upon request.

## Code availability

The code for the progression model is deposited at https://github.com/yohanjohn/CTE

